# Unprecedented sighting of coral populations and associated fauna in the High Sedimentation Waters of Puerto Velero, Atlántico, Colombia

**DOI:** 10.1101/2024.10.09.617471

**Authors:** Jorge Moreno-Tilano, Irene Lucia Sanabria Ramirez, Rafik Neme

**Affiliations:** Department of Chemistry and Biology, Universidad del Norte, Barranquilla, Colombia; Max Planck Partner Group – Genomics and Biodiversity of the Colombian Caribbean; Colegio Alemán de Barranquilla– Deutsche Schule Barranquilla, Barranquilla, Colombia

**Keywords:** Anthropogenic impact, turbid corals, Colombian Caribbean

## Abstract

The marine coastal zone of Atlántico, Colombia, faces significant anthropogenic impact due to sediment and organic matter from the Magdalena River. This study reports the presence of two distinct coral patches in Puerto Velero, Atlántico, an area previously considered unfavorable for coral growth. Located at depths of 1-6 meters, these patches harbor a variety of marine species, extending the known distribution of 15 species in the Colombian Caribbean. Despite high sedimentation rates, the larger coral colonies show resilience, surviving mass mortality events and diseases. These findings highlight the adaptive capacity of corals in adverse conditions, emphasizing the importance of systematic monitoring and conservation efforts to understand their response to environmental changes and support marine biodiversity in the Colombian Caribbean.

## Introduction

The marine coastal zone of the Atlántico department represents the area of the Colombian Caribbean coast with the highest anthropogenic impact. This phenomenon is mainly attributed to the deposition of sediments and dissolved organic matter from the mouth of the Magdalena River, the country’s most important river in extension and benefited population, with a mean suspended sediment load of 142 million tons per year (Restrepo et al. 2017). Consequently, the oceanographic conditions of this region are extremely adverse for the establishment of marine species, particularly for sessile organisms, whose survival critically depends on the stability of their environment.

Corals in the Caribbean and the Atlántico department are of particular interest due to the high turbidity caused by the river’s influence (Garzón-Ferreira and Díaz 2003). There are no official reports of the presence of corals in the department of Atlántico, although it is located between two departments where the presence of well-developed coral areas has been documented (INVEMAR-MINAMBIENTE 2020). however, there have been anecdotal reports from local residents about the presence of corals in areas of the department. Approximately 50 years ago, in the municipality of Tubará, a coastal spit formation occurred, currently known as the Puerto Velero spit (Anfuso et al. 2015). This geomorphological formation acts as a natural barrier, mitigating the direct impact of waves and reducing the entry of sediments and pollutants from the northern mouth. As a result, a bay-type environment has been generated, where oceanographic conditions differ notably from those prevalent in the rest of the department.

Previous reports have noted the presence of corals and fauna commonly associated with them on the pillars of a port in the area, which function as artificial substrate for the settlement and growth of corals (Durán-Fuentes et al. 2018, 2021; Gracia. et al. 2021; Carvajal-Floria and Gracia 2022; Contreras-Rueda et al. 2023; Moreno-Tilano et al. 2023). This study aims to document the presence of corals adapted to the environmental conditions imposed by the Magdalena River in natural substrates of the Department.

## Methods

### Study Area

The following study was conducted in the marine-coastal zone near the Puerto Velero spit, located in the department of Atlántico, Colombia (Figure 1). This area is characterized by variable sedimentation rates, which range from 10.5 5 mg cm^−2^ d^−1^ during the rainy season, influenced by currents coming from the Darien to the northeast, and up to 83.6 5 mg cm^−2^ d^−1^ in the dry season, driven by Caribbean currents, northeast winds and associated waves that generate a coastal current in a southwest direction, facilitating the transport of sediments from the Magdalena fluvial plume into the bay (Rangel-Buitrago et al. 2016).

**Fig 1.**
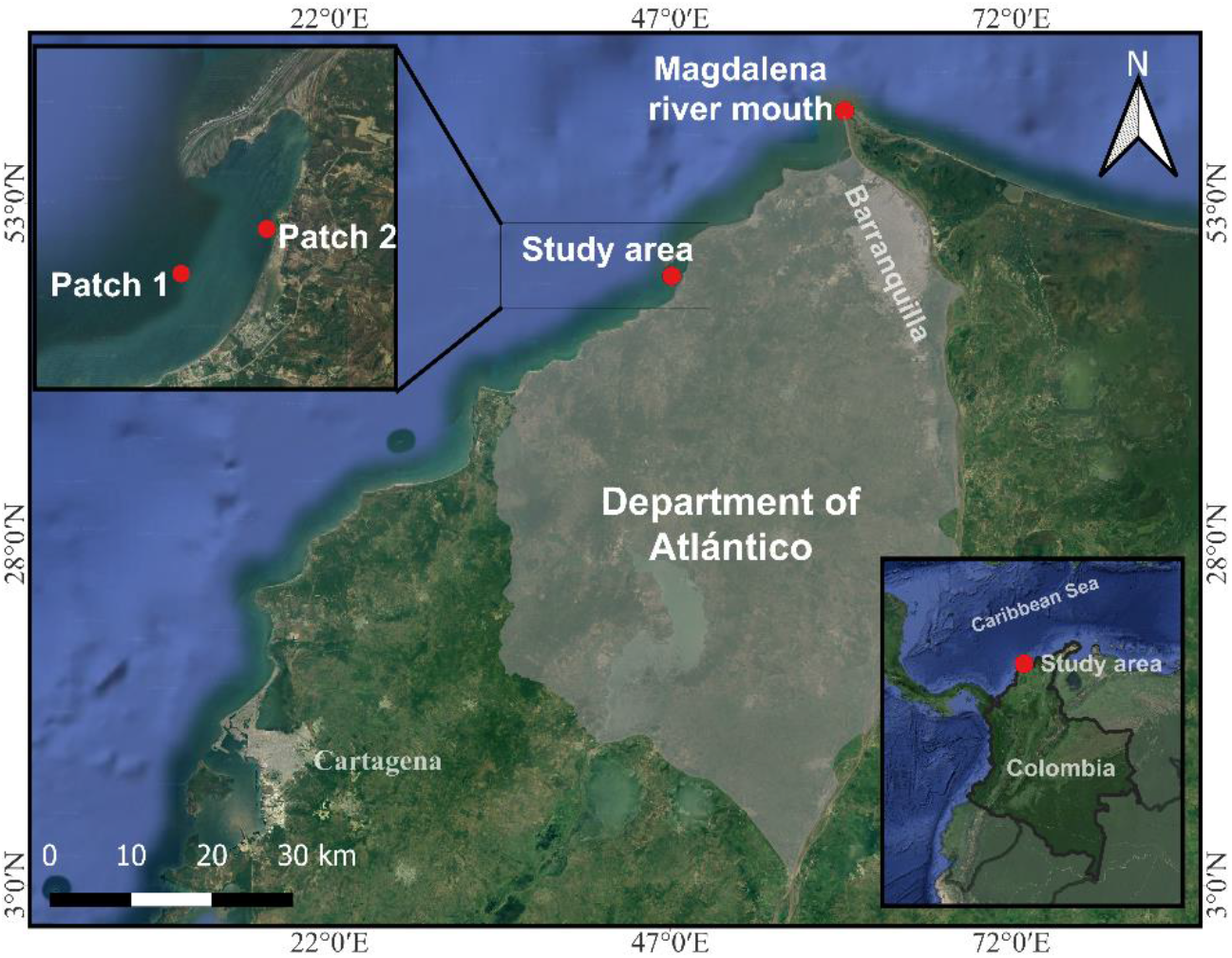
Map of the study area in the department of Atlántico, Colombia, and points of interest. “Patch 1” and “Patch 2” indicate the locations of the identified coral patches and where data were taken.

The average surface water temperature in the area is 28 °C, with an average salinity of 29.3 (Gracia et al. 2021). Climatic conditions in the area are determined by the north-south movements of the Intertropical Convergence Zone (ITCZ), giving rise to three distinct seasons: a dry season from December to April, a rainy season from August to November, and a transition season from May to July (Rangel-Buitrago et al. 2016).

The choice of this area for research was motivated by anecdotal records provided by the local fishing community, as well as recent evidence suggesting the presence of fauna commonly associated with coral communities in the region.

### Data Collection

The field exploration day took place on July 20, 2024. During the field exploration day, dives were conducted in random transects delimited within the study area. Data collection was based on video recording by means of a GoPro HERO7 Black camera, which was done in those sites where coral structures or associated fauna were identified. This method allowed the presence of organisms and structures of interest to be documented in detail in videos, thus ensuring that the data collection was as accurate and representative as possible of the biological diversity residing in the area.

### Video processing

The videos recorded during the dives were thoroughly reviewed in their entirety, playing them back at a reduced speed of 0.5x and performing a total of five complete reviews. This detailed approach allowed accurate identification of biological organisms present in the study area, including corals, fish, sponges and other invertebrates.

During the identification process, images of each organism were extracted from as many angles as possible, facilitating their subsequent taxonomic identification, which was carried out using taxonomic identification guides, such as those of Robertson et al. (2023), Reyes et al. (2002) and Humann (1993). These guides provided the necessary basis for classifying organisms to the most detailed taxonomic level possible and generating a list of taxa observed on our dives.

### Results and Discussion

We found two highly developed and compositionally distinct coral patches, each with an approximate area of 20 m^2^. The first patch, located at 10°54’52.84”N, 75°3’21.00”W at a depth of approximately 4 to 6 meters, is characterized by massive coral colonies with diameters exceeding 2 meters, predominantly of the species *Siderastrea siderea* (Fig. 2a) and *Porites* sp. (Fig. 2b).

**Fig 2.**
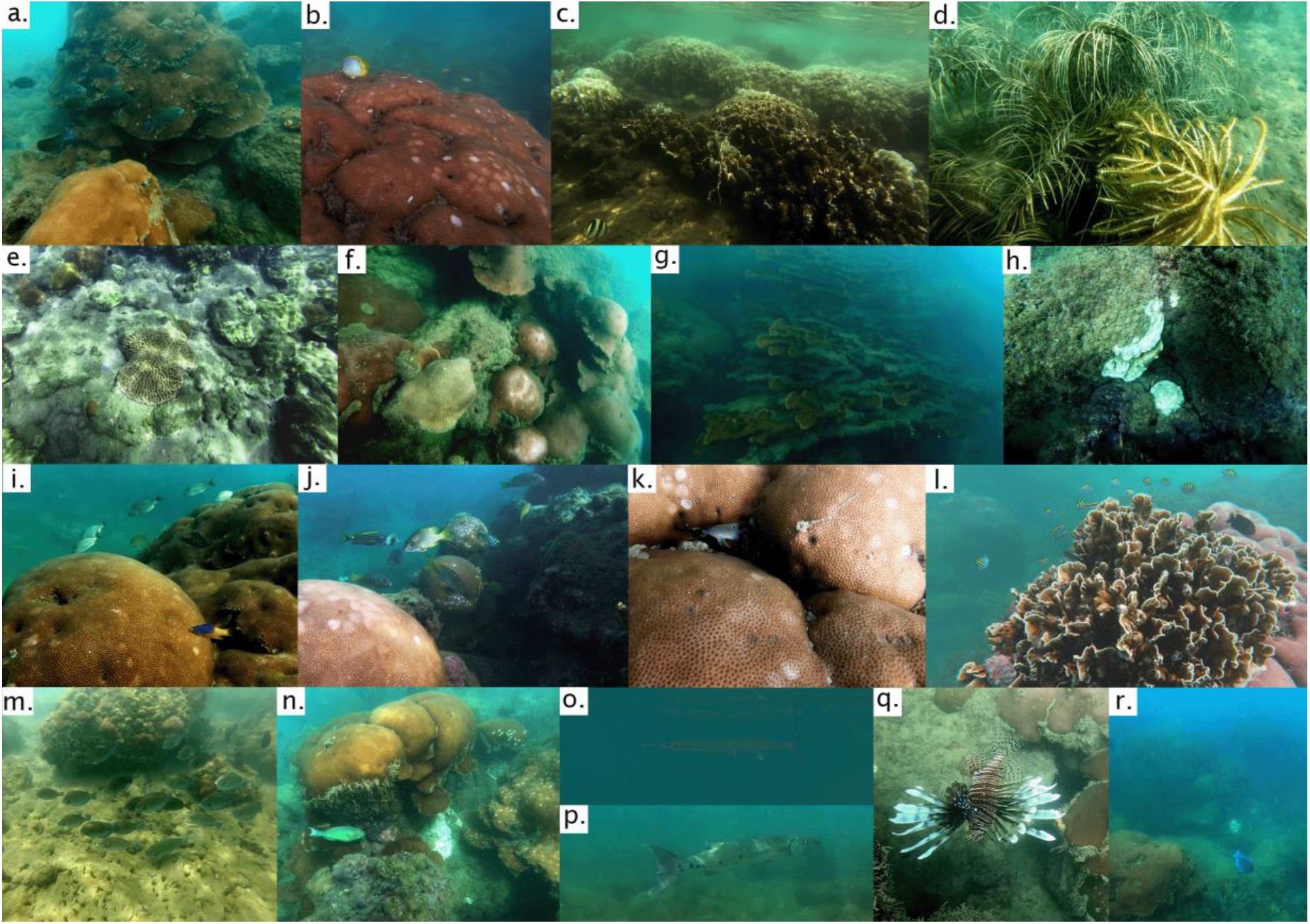
Photographic records for some species observed in two coral patches in Puerto Velero, Atlántico. **a**. *Siderastrea siderea, Orbicella sp*., *Acathusus tractus* and *Sparisoma rubripinne* (juvenile). **b**. *Chaetodon ocellatus* and *Porites sp*. (disease). **c**. *Chaetodon stristus* and *Millepora squarrosa* (bleaching). **d**. Octocorals. e. *Pseudodiploria sp*. **f**. *Faviidae sp*. **g**. *Acropora palmata* (healthy and dead specimens covered by algae). **h**. *Agaricia sp*. (bleaching). **i**. *Haemulon atlanticus, Bodianus rufus* and *Lutjanus apodus*. **j**. H*aemulon macrostoma, Abudefduf saxatilis, Lutjanus apodus, Sparisoma rubripinne* (juvenile) and *Porites sp*. (disease). **k**. *Stegastes adustus* (juvenile). **l**. *Elacatinus illecebrosus, Stegastes sp*., *Abudefduf saxatilis, Millepora squarrosa* and *Porites sp*. **m**. *Acanthurus tractus*. **n**. *Sparisoma rubiprinne*. **o**. *Tylosurus crocodilus* **P**. *Sphyraena barracuda*. **q**. *Pterois volitans*. **r**. *Acanthurus coeruleus*.

The second patch, located at 10°55’29.66”N, 75°2’10.49”W, 2.3 km from the first and at a depth of approximately 1 to 2 meters, is dominated by colonies of *Millepora squarrosa* (Fig. 2c) and various species of octocorals (Fig. 2d). These coral formations currently harbor diverse populations, including fish, polychaetes, echinoderms, and sponges, as captured in the videos (<u>Online resource 1, Online resource 2</u>). In total, 35 distinct taxa were identified across these groups, highlighting the rich biodiversity supported by this habitat (Table 1).

**Table 1.**
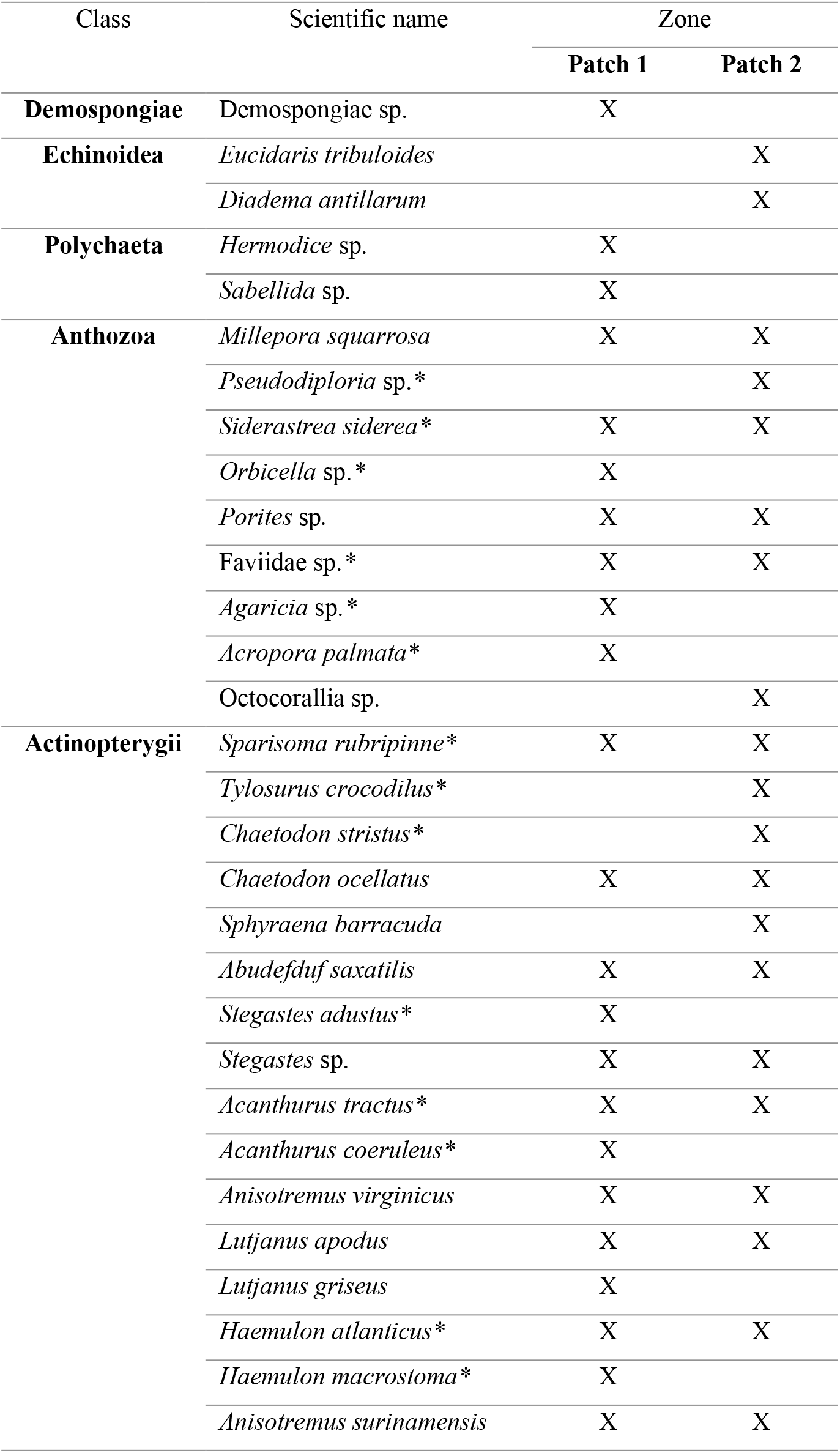
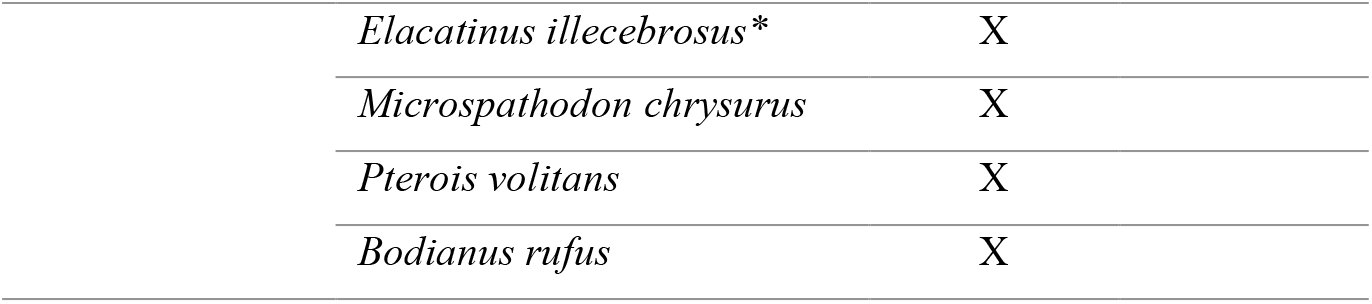
Taxa recorded on video in coral patches in Puerto Velero, Atlántico. The first column indicates the taxonomic class, the second column the scientific name of the taxon. The third and fourth columns indicate the presence of the taxon in Patch 1 and Patch 2, respectively. * Indicates species observed for the first time in the department of Atlántico.

| Class | Scientific name | Zone |  |
| --- | --- | --- | --- |
|  |  | Patch 1 | Patch 2 |
| **Demospongiae** | *Demospongiae* sp. | X |  |
| **Echinoidea** | *Eucidaris tribuloides* |  | X |
|  | *Diadema antillarum* |  | X |
| **Polychaeta** | *Hermodice* sp. | X |  |
|  | *Sabellida* sp. | X |  |
| **Anthozoa** | *Millepora squarrosa* | X | X |
|  | *Pseudodiploria* sp.* |  | X |
|  | *Siderastrea siderea* * | X | X |
|  | *Orbicella* sp.* | X |  |
|  | *Porites* sp. | X | X |
|  | *Faviidae* sp.* | X | X |
|  | *Agaricia* sp.* | X |  |
|  | *Acropora palmata* * | X |  |
|  | *Octocorallia* sp. |  | X |
| **Actinopterygii** | *Sparisoma rubripinne* * | X | X |
|  | *Tylosurus crocodilus* * |  | X |
|  | *Chaetodon stritus* * |  | X |
|  | *Chaetodon ocellatus* | X | X |
|  | *Sphyraena barracuda* |  | X |
|  | *Abudefduf saxatilis* | X | X |
|  | *Stegastes adustus* * | X |  |
|  | *Stegastes* sp. | X | X |
|  | *Acanthurus tractus* * | X | X |
|  | *Acanthurus coeruleus* * | X |  |
|  | *Anisotremus virginicus* | X | X |
|  | *Lutjanus apodus* | X | X |
|  | *Lutjanus griseus* | X |  |
|  | *Haemulon atlanticus* * | X | X |
|  | *Haemulon macrostoma* * | X |  |
|  | *Anisotremus surinamensis* | X | X |

|  |  |
| --- | --- |
| *Elacatinus illecebrosus\** | X |
| *Microspathodon chrysurus* | X |
| *Pterois volitans* | X |
| *Bodianus rufus* | X |

Notably, 15 of these species are not found in the lists of species reported for the department of Atlántico, thereby extending their known distribution within the Colombian Caribbean. The presence of these organisms, which collectively form a highly functional and continuously developing habitat, is particularly remarkable given the local environmental conditions. The discovery is especially surprising considering the high sedimentation rates in the area, ranging from 10.5 mg cm^−2^ d^−1^ to 83.6 mg cm^−2^ d^−1^ (Gracia et al. 2021), conditions that are generally unfavorable for species like corals, polychaetes, and sponges, which typically thrive in environments with lower turbidity(Golbuu et al. 2011). This makes their presence in this specific zone even more noteworthy.

We observed that the larger colonies have survived mass mortality events. Numerous individuals were colonized by macroalgae (Fig. 2g), while others exhibited partial or total bleaching processes (Fig. 2c, 2h). We also observed a significant prevalence of *Porites* specimens affected by ulcerative white spot disease (Figure 2b, 2i, 2j, 2k), a pathology documented as one of the main afflictions impacting corals in the Colombian Caribbean (Gil-Agudelo et al. 2009).

The presence and persistence of these coral formations, along with those recently documented in Varadero, Cartagena (López-Victoria et al. 2015; Pizarro et al. 2017), demonstrate a remarkable resilience and adaptation of the corals in this region of the Colombian Caribbean to the adverse environmental conditions generated by the continental discharges of the Magdalena River. The lack of previous reports on coral reefs in this area of tourist interest highlights the need to strengthen marine research in the Atlántico department, itself the population and research hub of the Colombian Caribbean in other areas of knowledge. Consequently, the characterization, study, and systematic monitoring of these coral ecosystems are of vital importance, both for their conservation and to understand their response to global environmental changes associated with climate change.

## Acknowledgements

We gratefully acknowledge the funding provided by the Max Planck Society and Universidad del Norte, which made this research possible. We also thank the reviewers for their valuable comments and suggestions

## Author Contributions

All authors contributed to the conception and design of the study. Material preparation, data collection and analysis were performed by Jorge Moreno-Tilano, Irene Lucia Sanabria Ramírez, and Rafik Neme. The first draft of the manuscript was written by Jorge Moreno-Tilano and all authors commented on previous versions of the manuscript, then the final version was done by Rafik Neme. All authors read and approved the final manuscript.

## Data Availability

The videos obtained and analyzed in this study can be found in the following hyperlinks: Online resource 1, Online resource 2

